# Heparins enhance C1 esterase inhibitor activity: a promising remedy for acute hereditary angioedema

**DOI:** 10.1101/2022.02.10.479965

**Authors:** Yingyan Zhou, Abraham Majluf-Cruz, Jaclyn Dennis, Eric Woroch, Lilian Hor, Brandon Hellbusch, Erin Archuleta, Lorelenn Fornis, Cindy Garcia, Shanae L. Aerts, Xiyuan Bai, Shaun Bevers, Eric P. Schmidt, Melanie Bates, Randolph V. Fugit, Sandra Nieto-Martinez, Manuel Galvan, Patricia Giclas, Ashley Frazer-Abel, Edward D. Chan

## Abstract

**Rationale:** Hereditary angioedema (HAE) is a potentially life-threatening illness most commonly due to deficiency or dysfunction of C1-esterase inhibitor (C1-INH). While specific treatments are available to thwart acute exacerbations, they are extremely costly and some can be associated with rare but serious side effects. The heparins are long known to augment C1-INH activity and case reports / series have documented their efficacy in treating HAE.

**Objective:** to determine if unfractionated heparin and two low-molecular weight heparins (enoxaparin and nadroparin) can augment C1-INH activity *ex vivo* in the sera of patients with HAE and in an *in vitro* biochemical assay.

**Methods:** C1-INH activity in the absence or presence of the heparin formulations were analyzed by two different methods. To measure C1-INH activity *ex vivo*, a commercially available assay was utilized with patient sera, excess amounts of C1s, and a substrate of C1s which, upon cleavage by C1s, produces a chromogenic product. To determine biochemically the C1-INH activity *in vitro*, a pharmacologic grade C1-INH, recombinant C1s (C1s-CCP12SP), and a peptide substrate of C1s were employed. Microscale thermophoresis was used to determine whether C1-INH binds to heparin.

**Main results:** in patient sera, nadroparin was superior to enoxaparin and unfractionated heparin in augmenting C1-INH activity, followed by enoxaparin and then unfractionated heparin. In the *in vitro* biochemical assay, all three heparins augmented C1-INH-C1s binding linearly in a dose-dependent fashion. Microscale thermophoresis assay demonstrated that nadroparin binds to C1-INH, providing a mechanism by which heparin facilitates the interaction between C1-INH and the proteases known to produce bradykinin, the mediator of HAE.

**Conclusion:** low-molecular weight heparin augments C1-INH activity and should be studied as a potential treatment for acute HAE.

## INTRODUCTION

Hereditary angioedema (HAE) is an autosomal dominant disorder caused by deficiency or dysfunction of C1 esterase inhibitor (C1-INH, gene *SERPING1*) – known as Type 1 HAE (~85% of cases) and Type 2 HAE, respectively. This lack of functional C1-INH results in decreased inhibition of kallikrein and plasmin, serine proteases that converts high-molecular weight kininogen to bradykinin (1). Binding of excessive amounts of bradykinin to its B2 receptor on endothelial cells causes vasodilation and increased permeability of the post-capillary venules, culminating in capillary leak that characterizes HAE. Bradykinin can also directly degranulate serosal mast cells of the rat and hamsters but not of humans (2). Approximately 25% of Type 1 and 2 HAE cases result from spontaneous gene mutations with no family history. Another rare type of HAE (Type 3) is associated with normal C1-INH level and function, is estrogen-dependent, and thus more common in females (3); some cases are due to a missense gain-of-function mutation of coagulation Factor XII gene. Other rarer forms of Type 3 HAE include loss-of-function mutation of angiopoietin-1 (which normally antagonizes the vascular permeability of bradykinin) (4) or gain-of-function mutations of plasminogen (precursor of plasmin) (5) or kininogen-1 (precursor to high-molecular weight kininogen) (6). Furthermore, associated loss-of-function or gain-of-function mutations of other proteins in the Kallikrein-Kinin System (KKS) – such as carboxypeptidases (normally able to remove arginine residues at the carboxy-terminus of bradykinin), angiotensin converting enzyme 2 (normally inactivates bradykinin), neutral endopeptidases (normally deactivates kinin), and the bradykinin receptors – could modify the phenotypic presentations of Type 1 and 2 HAE (7).

Pharmacologic treatments for acute HAE attacks include C1-INH replacement (plasma-derived or recombinant), ecallatide (kallikrein inhibitor), and icatibant (a bradykinin receptor antagonist) (1, 8–10). These agents are effective but are very expensive and rarely associated with serious side effects such as anaphylaxis with C1-INH and ecallatide. Oral kallikrein inhibitors used prophylactically significantly lowered the attack rate for acute HAE (11) or shortened the angioedema episodes (12). Another kallikrein inhibitor lanadelumab administered subcutaneously for 26 weeks significantly reduced the number of acute exacerbations of HAE (13).

Nearly 100 years ago, heparin was reported to potentiate C1-INH activity in the classical complement pathway (14, 15). Hypothesized mechanisms include the ability of heparin to enhance binding of C1-INH to C1 esterase (C1s) and interference of C1q binding to an activator (15–17). While the complement pathway *per se* is not considered to play a pathogenic role in HAE, C1-INH also inhibits kallikrein and plasmin, the two key serine proteases that produce bradykinin. Thus, the ability of the heparins to augment C1-INH activity may be highly relevant in the context of preventing or treating acute attacks of HAE. Indeed, unfractionated heparin (UFH) and low-molecular weight heparins (LMWH) have shown efficacy in the treatment of acute HAE attacks in case reports and series (18–21). Whether heparins can augment C1-INH activity in the sera of HAE patients is unknown. Herein, we evaluated UFH and two formulations of LMWH on C1-INH activity with HAE patient sera as well as their ability to directly bind C1-INH.

## EXPERIMENTAL METHODS

### Materials

Unfractionated heparin sodium (1,000 units/mL) was obtained from Fresenius Kabi, Lake Zurich, IL. Enoxaparin sodium (100 mg/mL) was obtained from Winthrop, Bridgewater, NJ. Nadroparin (Fraxiparine, 950 IU of anti-Xa per 0.1 mL) was obtained from Aspen Holdings Pharmaceutical, Durban, South Africa. C1-INH (Berinert) used for the continuous kinetic assay was purchased from CSL Behring and further purified as previously described (16). C1-INH (1 mg/mL from normal human serum) used for the Microscale Thermophoresis assay was purchased from Complement Technology (Tyler, TX). Z-Lys-SBzl was purchased from Bachem. 4,4’-dithiodidyridine (DTDP) and heparin sodium salt (porcine intestinal mucosa) were purchased from Sigma-Aldrich. C1s-CCP12SP was expressed, refolded and purified as previously described (22). Microscale Thermophoresis (MST) was performed with a Monolith NT.115 Pico instrument (Nanotemper Technologies, Munchen, Germany). Monolith Protein Labeling Kit RED-NHS 2nd Generation (Amine Reactive) and Monolith NT.115 Capillaries, used in the MST experiments, were purchased from Nanotemper Technologies.

### Serum samples of patients with known HAE

Twenty-seven de-identified serum samples of patients with known HAE were obtained from Biobank Resources at National Jewish Health (NJH) through the Honest Broker Protocol.

### Determination of C1-INH level

C1-INH protein levels were measured using the SPAplus automated protein analyzer (The Binding Site Group Ltd, Birmingham, UK) – which utilizes turbidimetric measurement to determine protein level – according to manufacturer’s instructions.

### Concentrations of UFH, enoxaparin, and nadroparin tested

A standard initial dose of UFH given to patients is 5,000 units subcutaneously or intravenously for prevention or treatment of venous blood clots. Since the average blood volume is approximately five liters and the hematocrit is ~45%, the plasma volume is 2.75 or approximately 3 liters. Assuming UFH is given intravenously, the concentration is 5,000 units per 3000 mL = 1.7 U/mL. Thus, the range of UFH concentrations chosen to test the effect of UFH on C1-INH activity by Method 1 (see below) are 0, 0.25, 0.5, 1, 1.5, 2, 4, and 8 U/mL. For the standard enoxaparin dose of 1.5 mg/kg, calculations based on a 70 kg person and three liters of plasma gives an enoxaparin concentration of 35 μg/mL, assuming all the enoxaparin administered is absorbed into the circulation. Thus, the final enoxaparin concentrations used to determine its effects on C1-INH activity are 0, 8.8, 17.5, 35, 52.5, and 70 μg/mL. For nadroparin, with a 0.1 mL/kg dose of a 5,700 units per 0.6 mL solution, gives an estimated plasma concentration of 2.2 U/mL for a 70 kg person with 3 liters of plasma. Thus, the final nadroparin concentrations used to determine its effect on C1-INH activity are 0, 0.55, 1.1, 2.2, 3.3, 4.4, and 8.8 U/mL.

### Determination of C1-INH activity

#### Method 1. Chromogenic assay to measure C1-INH activity

C1-INH activity in the absence and presence of varying concentrations of heparins was measured in the 27 serum samples using a commercially available chromogenic assay (Technochrom C1-INH, Technoclone GmbH, Vienna, Austria), adapted from the manufacturer’s instructions. In this assay, serum – which contains C1-INH – is incubated with excess C1 esterase (C1s). C1s that is not bound to and inactivated by C1-INH is able to cleave a substrate (C_2_H_5_CO-Lys(e-Cbo)-Gly-Arg-pNA) that develops color (is chromogenic). Hence, increased C1-INH activity in serum results in greater inhibition of C1s, leading to lower chromogenic product. Data were analyzed using regression analysis from a reference standard curve. The samples were tested in duplicate wells.

#### Method 2. Continuous kinetic assay to measure C1-INH activity

The C1-INH inhibitor activity in the presence of each of the three clinical-grade heparin formulations was also measured biochemically (in the absence of serum) by an absorbance-based assay employing C1-INH (Berinert), C1s-CCP12SP, Z-Lys-SBzl as the substrate, and the chromogenic thiol reagent 4,4’-dithiodipyridine (DTDP) (23, 24). Residual C1s activity was monitored at 324 nm. Assays were conducted in 20 mM Tris, 250 mM NaCl, 0.005% Triton X-100, 5% dimethyl sulfoxide, pH 7.4 at 30°C in a FLUOstar Omega plate reader (BMG Labtech). Progress curves were fitted by non-linear regression using OriginPro 2018 (OriginLab Corporation) to the integrated rate equation for slow binding inhibition to determine the observed pseudo-first-order rate constant (16, 25, 26). Results generated in the presence of each of the heparin formulations are expressed as the fraction to that generated in the absence of heparin (fold potentiation).

### Microscale thermophoresis

Microscale thermophoresis (MST) was used to analyze the binding between C1-INH and nadroparin since we found nadroparin to be the most potent in augmenting C1-INH activity. C1-INH protein (Complement Technology) was fluorescently labeled according to the manufacturer’s instructions (Nanotemper Technologies). The nadroparin stock concentration was serially diluted two-fold in nano-pure water to generate 16 different concentrations (1.28×10^−7^ to 4.2×10^−3^ M). Next, fluorescently labeled C1-INH in PBS buffer + 0.05% P20 was added such that the final concentration of C1-INH in each tube was 10 nM. Samples were then loaded into Monolith NT.115 Capillaries and analyzed. In terms of instrument settings, a Pico.RED detector was used at 20% laser power and medium MST power was used for all experiments. Data were analyzed using the MO Affinity Analysis software suite (Nanotemper Technologies).

### Statistical analyses

The significance of fold potentiation of C1-INH activities was calculated using two-way ANOVA and Dunnett’s multiple comparison test, using GraphPad Software, San Diego California USA.

## RESULTS

### C1-INH levels

Of the 27 serum samples from patients with known HAE, the serum concentration of C1-INH are shown in **Table 1**. As shown, 24 of the 27 samples had C1-INH levels below the lower limits of normal; the three samples within the range of normal C1-INH levels were at the lower end.

**Table 1.** C1-INH levels.

| **Serum Sample No.** | **C1-INH level (mg/dL)**<br>**(Normal range: 20-37**<br>**mg/dL)** |
| --- | --- |
| 1 | 8.0 |
| 2 | 21.0 |
| 3 | 11.0 |
| 4 | 7.0 |
| 5 | 9.0 |
| 6 | 9.0 |
| 7 | 7.0 |
| 8 | 23.0 |
| 9 | 12.0 |
| 10 | 9.0 |
| 11 | 10.0 |
| 12 | < 6.0 |
| 13 | 7.0 |
| 14 | 14.0 |
| 15 | < 6.0 |
| 16 | 13.0 |
| 17 | 7.0 |
| 18 | 7.0 |
| 19 | 8.0 |
| 20 | 10.0 |
| 21 | 6.0 |
| 22 | 11.0 |
| 23 | 7.0 |
| 24 | 14.0 |
| 25 | 8.0 |
| 26 | 7.0 |
| 27 | 21.0 |

### UFH, enoxaparin, and nadroparin differentially augmented C1-INH activity in sera from HAE patients

There was a slight increase in C1-INH activity with increasing UFH concentrations in most of the samples (**Fig 1A**). However, at the highest UFH concentration of 8 U/mL, a white precipitate formed in some samples, resulting in a spurious decrease in measured C1-INH activity in these samples (**Fig 1A**). The plot of the mean C1-INH activity as a function of the UFH concentration showed the dose-dependent increase in C1-INH activity, peaking at 4 U/mL UFH; indeed, the only UFH concentration in which the C1-INH activity was above the 40% of normal activity (laboratory normal range is 74% to 174%) – traditionally reported as a threshold level above which affords protection against symptomatic HAE (27–30) – was the 4 U/mL of UFH (**Fig 1B**).

**Figure 1.**
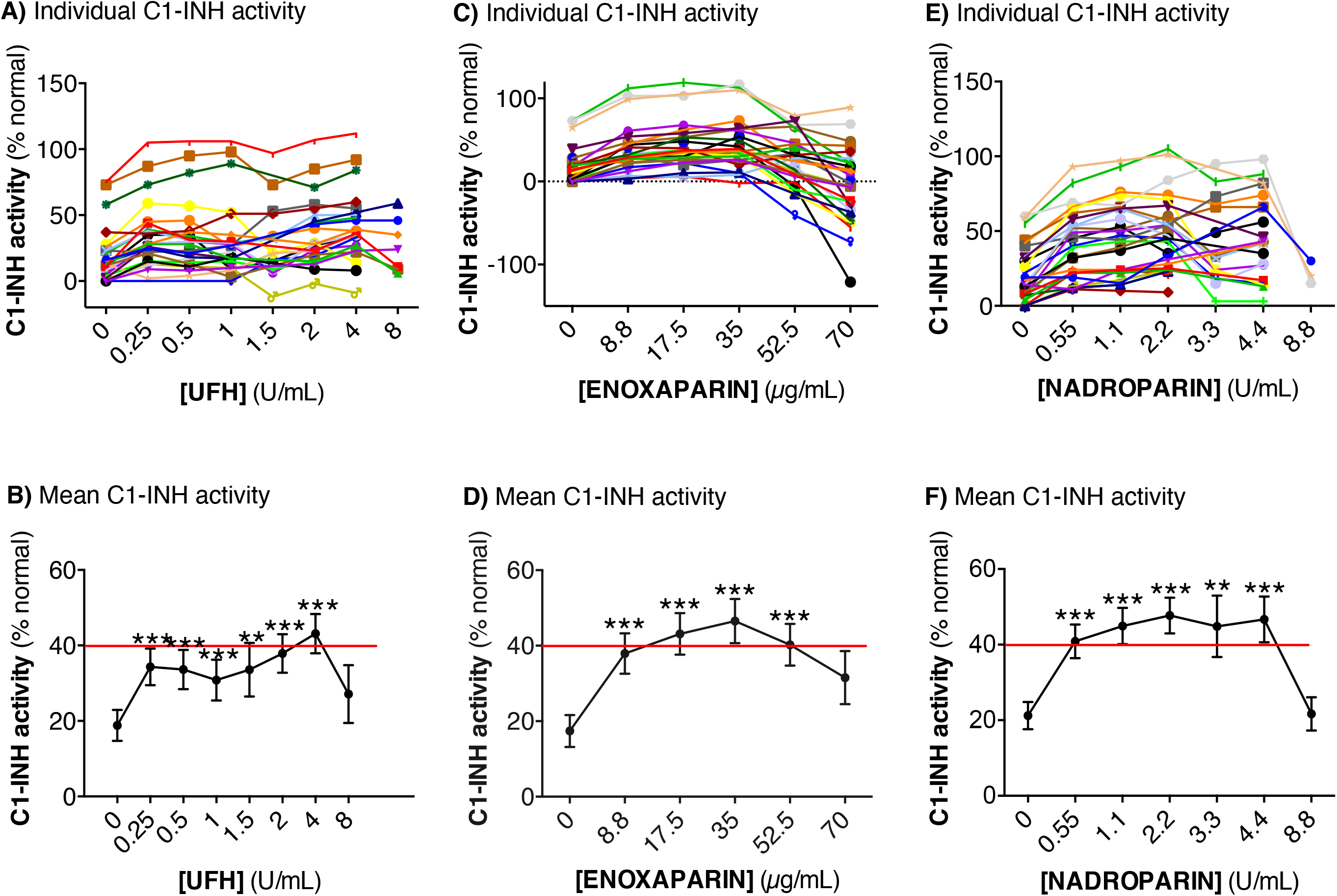
Effects of the heparins on C1-INH activity using sera of patients with hereditary angioedema. Effect of **(A/B)** clinical-grade UFH, **(C/D)** enoxaparin, and **(E/F)** nadroparin on C1-INH activity in sera of 27 patients with hereditary angioedema. Data are reported as the effects of each of the heparins on the C1-INH activity of the individual sera (**A/C/E**) and the mean results (**B/D/F**). **p<0.01, ***p<0.001 compared with zero amount of heparin formulations added.

Incubation of the HAE sera with enoxaparin demonstrated an increase in C1-INH activity with the lowest enoxaparin concentration tested, followed by a steadier rise (**Fig 1C**). Similar to that seen with UFH, precipitation of the mixtures occurred at the higher enoxaparin concentrations, resulting in a spurious decrease in C1-INH activity (**Fig 1C**). For the mean level of C1-INH activity as a function of enoxaparin concentrations, the peak C1-INH activity occurred at an enoxaparin concentration of 35 μg/mL (**Fig 1D**).

The effect of nadroparin on increasing C1-INH activity was significantly greater than for either UFH or enoxaparin, with an increase in C1-INH activity above the 40% cut-off line even with the lowest nadroparin concentration used (**Fig 1E / F**). Hence, there were significantly more nadroparin concentrations tested that increased the mean C1-INH activity above the 40% of normal activity than that for either UFH or enoxaparin (compare Fig 1F with 1B and 1D). Precipitation also occurred with the highest concentrations of nadroparin in couple of the samples that were tested, resulting in spuriously low C1-INH activity (**Fig 1E / F**).

### Increase in C1-INH activity to ≥40% as a function of baseline C1-INH level

In the absence of serum, UFH, enoxaparin, or nadroparin had zero C1-INH activity as measured by the chromogenic assay (data not shown). Thus, we determined whether the increase in C1-INH activity by the heparins was dependent on the baseline concentration of C1-INH in the sera. For this analysis, we first categorized each of the 27 serum samples on whether there was an increase in C1-INH activity to >40% of normal C1-INH activity with each of the heparins at any concentration tested. We then plotted these binary categories as a function of the baseline C1-INH concentrations. Such analysis showed that in the samples in which C1-INH activity increased to >40% of normal activity with the addition of UFH or enoxaparin, the mean C1-INH level trended to be lower (**Fig 2A/B**). However, in patient serum samples in which addition of nadroparin increased C1-INH activity to >40% of normal, the mean plasma C1-INH activity was significantly lower than those serum samples in which there was no increase in C1-INH activity above the 40% cut-off (**Fig 2C**). These findings further support the notion that the heparins, especially nadroparin, are able to significantly augment C1-INH activity in serum samples with low C1-INH levels are low, as seen in Type 1 HAE.

**Figure 2.**
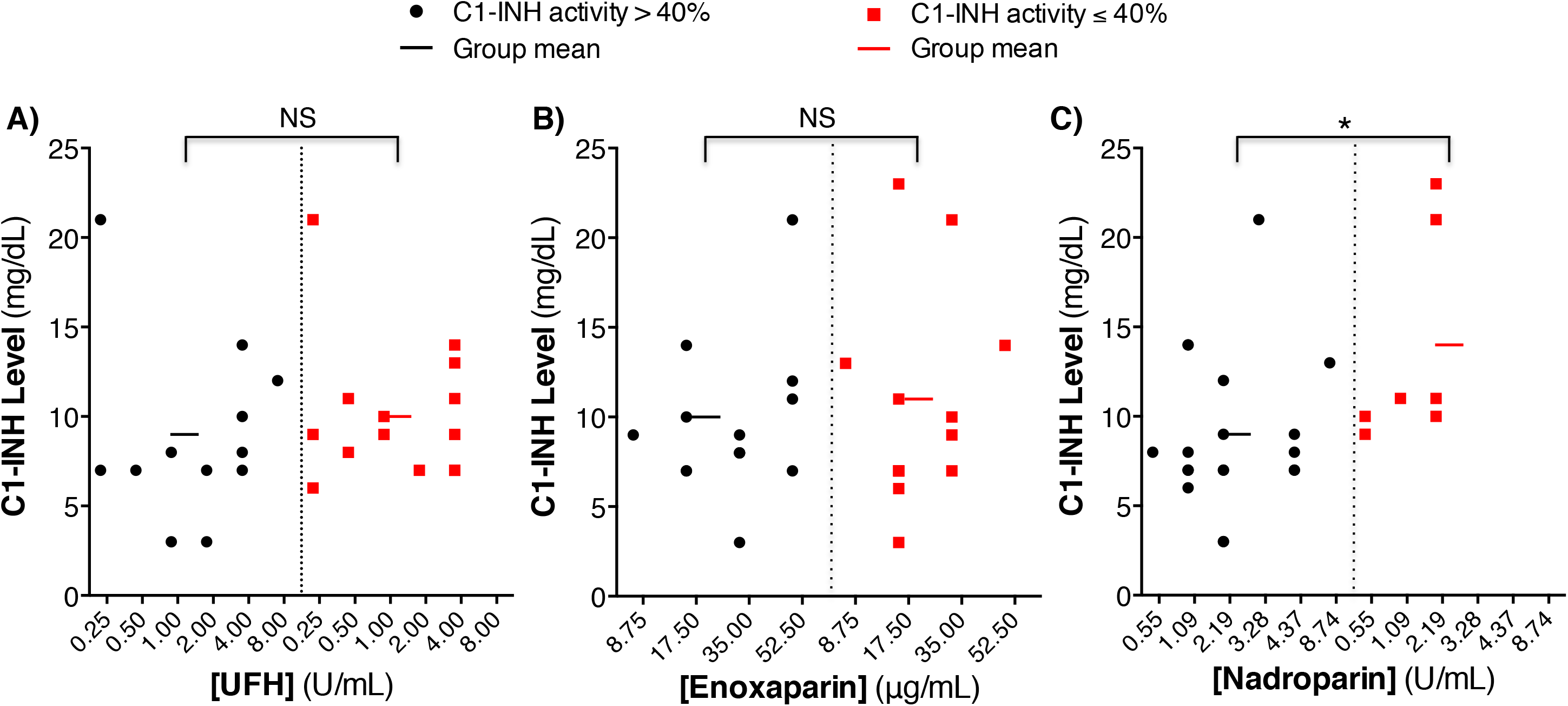
Analyzing the effects of the heparins in increasing C1-INH activity to >40% activity as a function of the baseline C1-INH level. The mean C1-INH level was stratified between those with any increase in C1-INH activity >40% above baseline vs. those that did not have such an increase (≤40% of baseline) in the presence of each of the three clinical-grade heparin formulations. *p<0.05.

### UFH, enoxaparin, or nadroparin increases C1-INH activity in the continuous kinetic assay

Results in the serum-based chromogenic assay were confirmed with an alternative approach to rule out the effect of serum proteins in the measurement of C1-INH activity. In this assay, commercially available C1-INH (Berinert) was incubated with a recombinant form of C1s and a peptide substrate for C1s (Z-Lys-SBzl) was added alone or with increasing concentrations of the each of the three heparins administered to patients. We found that all three heparin preparations used clinically significantly potentiated C1-INH-C1s interaction to a similar extent, resulting in faster inhibition of C1s with less chromogenic product being produced (**Fig 3A/B/C**). The vertical dashed lines in three of the graphs in Fig 3 correspond to the highest concentrations of the three heparins used in Fig 1, showing that the augmentation of C1-INH activity with heparin concentrations which are more in the physiologic range has a direct linear relationship.

**Figure 3.**
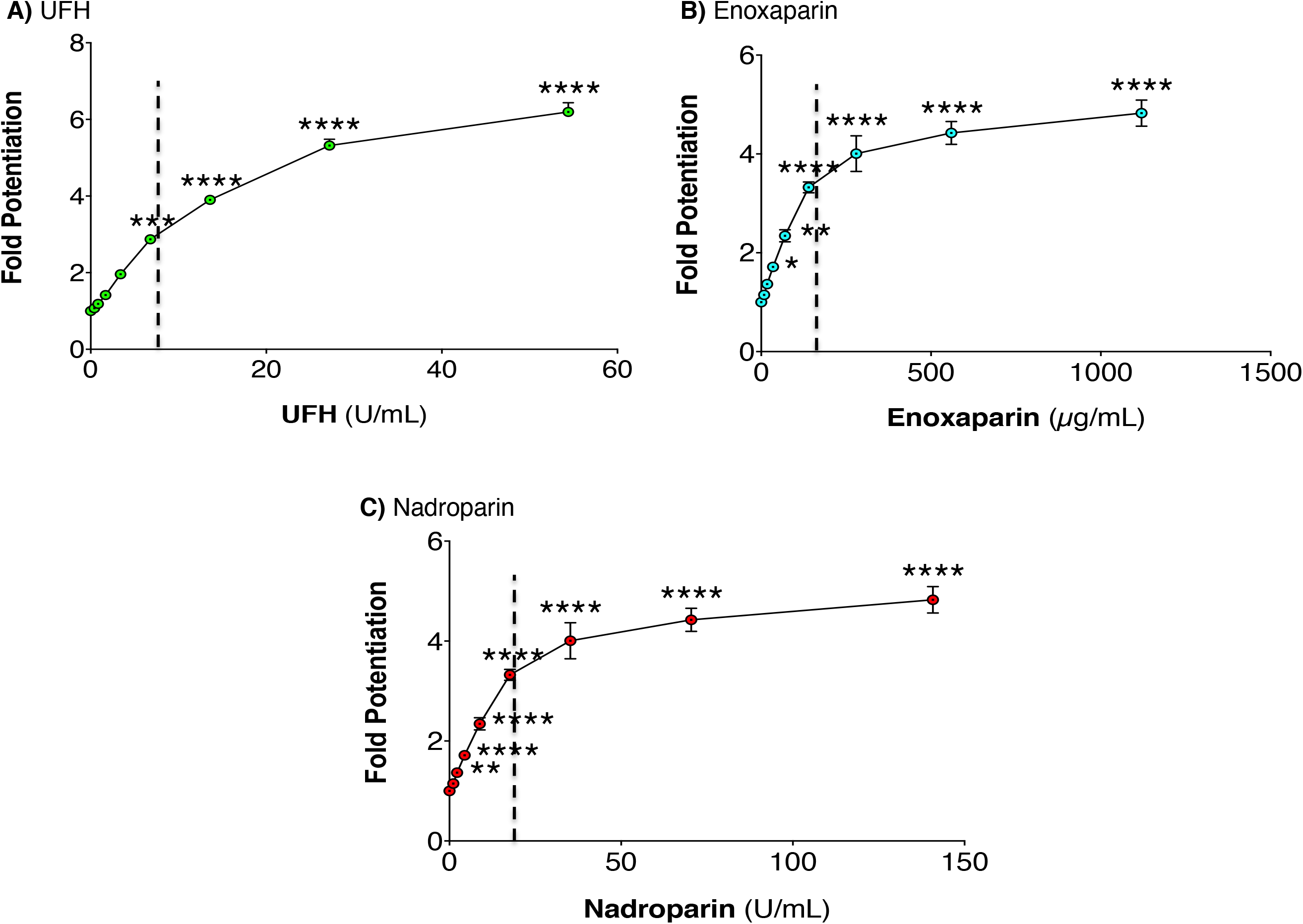
Effect of heparin on the observed rate of association of C1-INH and C1s. The effect of clinical-grade **(A)** UFH, **(B)** enoxaparin, **(C)** nadroparin or **(D)** research-grade UFH on the observed rate of association between recombinant C1s and C1-INH was determined at increasing concentrations. Fold potentiation, the ratio of the pseudo-first-order rate constant (kobs) in the presence of heparin to that in the absence of a cofactor, is plotted as a function of heparin concentrations. *p<0.05, **p<0.01, ***p<0.001, ****p<0.0001 compared with zero amount of heparin formulations added.

### Nadroparin binds to C1-INH as measured by Microscale Thermophoresis

Since nadroparin was the most potent in augmenting C1-INH activity in the sera of HAE patients, Microscale Thermophoresis (MST) was employed to determine whether nadroparin binds to the C1-INH protein. MST measures the mobility of a fluorescent labeled protein in the presence of a thermal gradient, a process known as thermophoresis. The thermophoretic mobility of a molecule is dependent on a variety of factors including charge, hydrodynamic radius and solvent. If the fluorescent labeled molecule is exposed to another molecule to which it binds, the thermophoretic mobility of that fluorescent labeled molecule will change in a measurable amount. For this experiment, 10 nM of fluorescent labeled C1-INH was combined with 16 different nadroparin concentrations (1.28×10^−7^ to 4.2×10^−3^ M) placed in 16 different capillary tubes, after which the thermophoretic mobility of the C1-INH was measured by fluorescence. The thermophoretic mobility of the fluorescent-labeled C1-INH changed in a concentration-dependent manner with increasing concentrations of the nadroparin in the mixture (**Fig 4**), indicating that C1-INH binds to nadroparin with a K_d_ of 3.34 × 10^−4^ M.

**Figure 4.**
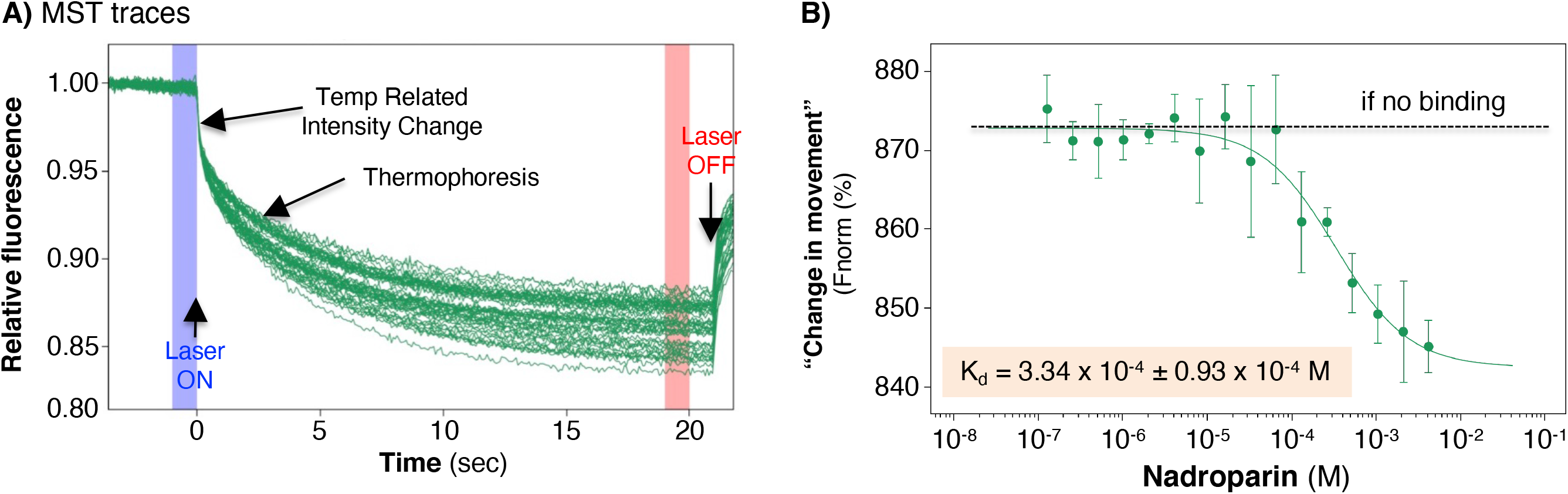
Microscale thermophoresis to determine the presence of binding between nadroparin and C1-INH. **(A)** Microscale thermophoresis of the time-course tracings following the mixing of 10 nM of fluorescent-tagged C1-INH with 16 different nadroparin concentrations (1.28×10^−7^ to 4.2×10^−3^ M) in capillary tubes and subjected to a thermogradient. Heat is applied to the capillary tubes by a laser beam to create a thermal gradient and heat turned off. **(B)** The presence of a “change in movement” of the fluorescent-tagged C1-INH with varying nadroparin concentrations demonstrates that the two molecules interacted *in vitro*. Data shown are the mean and SD of three independent experiments.

## DISCUSSION

HAE is characterized by recurrent episodes of non-pruritic, non-pitting, subcutaneous, and submucosal edema typically involving the arms, legs, hands, feet, bowels, genitalia, trunk, tongue, face, and/or larynx (8). The edema typically increases slowly over 24 hours and gradually subsides over the next 48 to 72 hours. Foremost in the treatment of angioedema is ensuring that the upper airway does not become compromised or if obstruction is already present, immediate steps are taken to secure a patent airway. For true HAE, epinephrine, glucocorticoids, and anti-histamines are ineffective. More specific acute HAE treatment include one of several C1 inhibitor concentrates given intravenously, or specific inhibitors of the contact pathway – ecallantide and icatibant – administered subcutaneously. The major side effect of icatibant is discomfort at injection site and the most serious one for C1-INH replacement therapies and ecallantide is anaphylaxis.

Heparin has been shown to inhibit several components of the complement pathway: *(i)* interferes with binding of C1q to immune complexes, *(ii)* inhibits the interaction of C1s with C4 and C2, *(iii)* inhibits C2 binding to C4b, and *(iv)* facilitates the binding of C1s and C1-INH to augment C1-INH activity (23, 31–33). Heparin is known to deplete C1, a property that may be shared by other highly charged polyanionic substances (34). Despite these effects of heparin on the complement pathway, the complement system *per se* is not considered to play a primary role in the pathogenesis of HAE. Poppelaars and colleagues (15) demonstrated that UFH and three LMWH (dalteparin, enoxaparin, and nadroparin) individually inhibited the functional activity of the classical, alternative, and lectin complement pathways. Interestingly, all forms of heparin individually also potentiated C1-INH in inhibiting the classical and lectin complement pathways but less consistently augmented C1-INH activity in the functional assay for the alternative pathway (15).

In the current study, we investigated the effects of UFH, enoxaparin, or nadroparin on C1-INH activity in the sera of HAE patients. We found that nadroparin had the greatest effect in enhancing C1-INH activity, followed by enoxaparin and then UFH. These findings support work from nearly 20 years ago that LMWH augmented C1-INH activity on C1 more than UFH albeit the experiments were performed biochemically without the use of patient serum samples (31). Interestingly, when the C1 substrate was in a fluid phase, neither UFH or LMWH were effective in augmenting C1-INH activity on C1 whereas if the C1 substrate was bound to red blood cells, both UFH and LMWH augmented C1-INH activity on C1 (31).

Not every serum sample had an increase in C1-INH activity with the heparins. Perhaps these serum samples contain adequate levels of other protease inhibitors, which is plausible since in the presence of other protease inhibitors, antithrombin-heparin co-factor is not a quantitatively important inactivator of kallikrein (35). Nevertheless, our findings support the proof-of-concept that the heparins – especially nadroparin – augment C1-INH activity as measured by the chromogenic assay in sera from HAE patients. The chromogenic assay is indispensable for determination of C1-INH function and a critical component of the HAE diagnostic panel because C1s function is exclusively linked to C1-INH activity only and no other inhibitor. Therefore, the enhance activity of C1-INH measured by the assay would be predicted to inhibit the production of bradykinin as well. Potential mechanisms by which UFH and LMWH augment C1-INH and anti-thrombin III activities – both of which can inhibit kallikrein – are diagrammed in **Fig 5** (35–37). Why the two LMWH enhanced C1-INH activity more than UFH remains to be determined. Although all heparins are anionic polysaccharides, the products are comprised of a mixture of different chain lengths of sulfated glycosaminoglycans. For example, whereas the average molecular weight of UFH is 15,000 Da, the range is wide – estimated to be 3 to ≥ 30 kDa (38). In contrast, LMWH is isolated from UFH by various methods of fractionation or deploymerization and although the average molecular weight of enoxaparin and nadroparin are much lower – estimated to be 4,300 and 4,500 Da – even the LMWH are comprised of glycosaminoglycan chains of varying lengths. Our findings suggest that the shorter chain heparins may have greater ability to enhance C1-INH activity.

**Figure 5.**
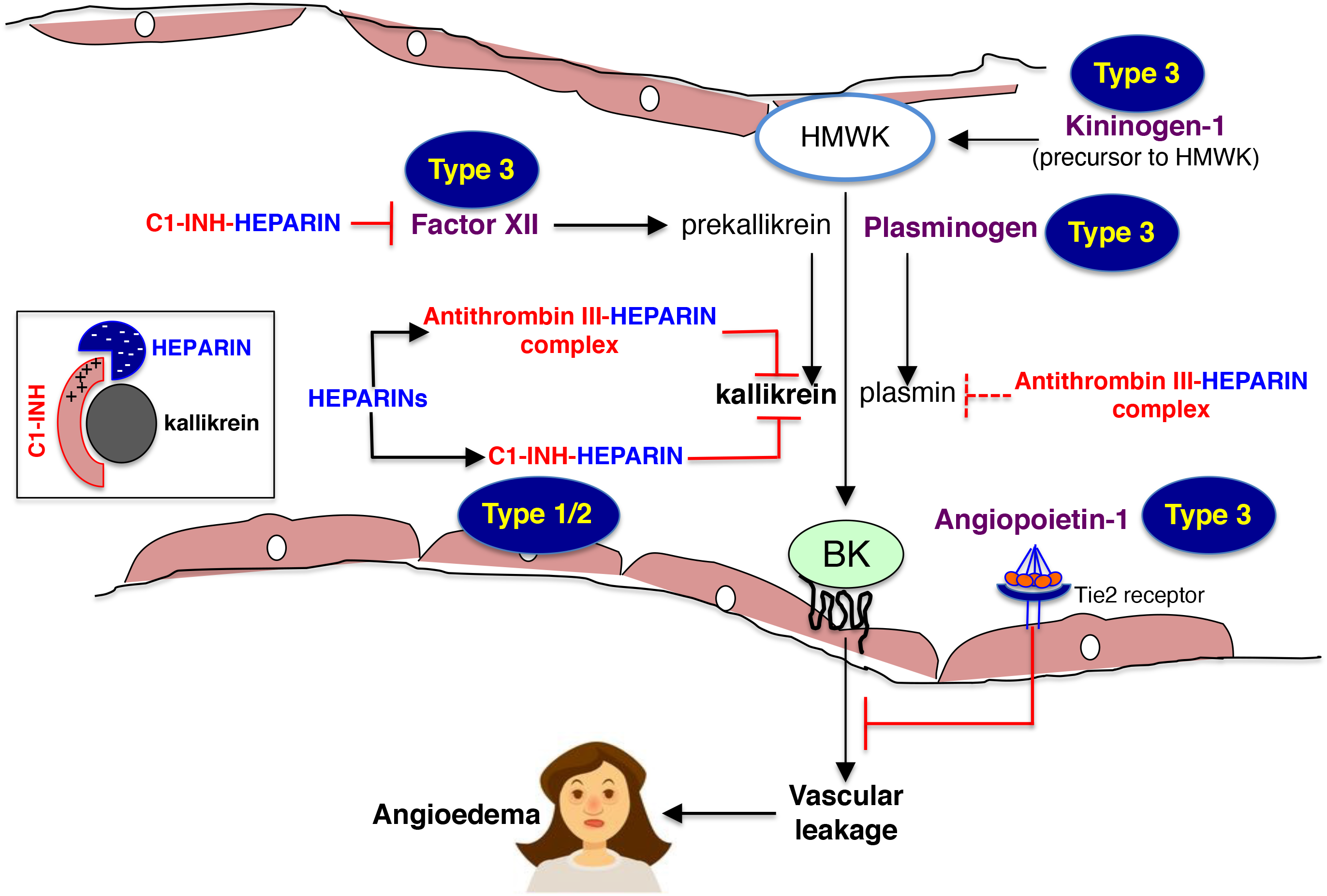
Diagram demonstrating how the heparins may inhibit the formation of bradykinin. For Types 1 and 2 HAE, heparins would be expected to bind C1-INH and antithrombin III, augmenting their activity to inhibit kallikrein activity and the subsequent formation of bradykinin. In addition, heparins could theoretically also antagonize the pathophysiologic processes that mediate the angioedema with normal C1-INH function (Type 3 HAE) due to mutation of plasminogen, angiopoietin-1, kininogen-1 (precursor to HMWK), or factor XII. Factor XII and HMWK are produced mainly by the liver but HMWK also produced by endothelial cells. Based on known activity of heparins to antagonize various parts of the pathways that lead the bradykinin formation, heparins could theoretically ameliorate all forms of HAE. **C1-INH**=C1 esterase inhibitor; **HAE**=hereditary angioedema; **HMWK**=high molecular weight kininogen.

We found that nadroparin bound to C1-INH, corroborating the finding of others that heparin binds C1-INH (31). This binding is highly significant since it appears that the ternary complex of heparin–C1-INH–protease is required for heparin to augment C1-INH inhibitory activity on the protease. Like other polysaccharides, heparins are negatively charged; indeed, heparin possesses the highest negative charge density of any known biological molecules. Furthermore, it is well known that polysaccharides such as dextran can potentiate C1-INH activity. More specifically, one negative charge dextran molecule was found to bind to multiple positively-charged F1 helix present in C1-INH, neutralizing some of the relevant positive charges on lysine and arginine residues on C1-INH. This interaction facilitates the binding of C1-INH to positively-charged autolysis loop of serine proteases, forming a ternary complex of polysaccharide–C1-INH–protease, and not “sandwiched” between the protease and anti-protease (C1-INH–polysaccharide–protease) (39). Extending this known effect of polysaccharide to our assay of C1-INH activity, we posit that the negatively-charged nadroparin and to a lesser extent enoxaparin and UFH also facilitated the formation of the heparin–C1-INH–C1s complex, potentiating the inhibitory effects of C1-INH on C1s. In the context of HAE, similar analogy may be made wherein heparin potentiates the inhibitory activity of C1-INH on kallikrein through formation of a heparin–C1-INH–kallikrein complex (**Fig 5, boxed diagram**). Other C1-INH-independent effects of the heparins in antagonizing bradykinin-mediated angioedema include the formation of the antithrombin–heparin complex that facilitates binding of antithrombin to both plasmin and kallikrein, thus inhibiting bradykinin formation (35–37).

Corroborating the biochemical evidence that the heparins augment C1-INH activity are published reports demonstrating the efficacy of heparin in treating HAE. In a randomized, double-blind, double-dummy, placebo-controlled, three-way crossover study comparing inhaled or subcutaneous heparin, subcutaneous heparin resulted in a significant decrease in flare intensity compared to inhaled heparin, and a trend toward decreased flare intensity compared to placebo (20). In 34 HAE patients with 256 attacks, nadroparin given subcutaneously after onset of prodromes and then every 8 to 12 hours for one day resulted in complete response in 90% (total cessation of an attack within two hours), partial response in 8%, and failure in 2% (19). Two patients with frequent attacks of HAE became “asymptomatic” when treated prophylactic doses of (weekly) nebulized heparin – up to 30,000 units per treatment (18); interestingly and paradoxically in one of the patient, the functional, plasma C1-INH level decreased after each heparin treatment.

The cost of all the HAE-specific therapeutics can be a barrier to use especially in low-income countries because the prices range from ~$3,000 to ~$15,000 per dose. In contrast, heparins are relatively inexpensive and have well-established safety profile; other than the obvious bleeding risk, heparin-induced thrombocytopenia is the most feared complication of the heparins but this risk is less with LMWH than UFH and can be easily monitored.

In summary, the results of this study using sera from HAE patients support the clinical reports of the beneficial effects of the heparins – especially LMWH – on acute symptoms of HAE. There are no randomized double-blind, placebo-controlled studies on the use of LMWH or its comparisons with the current FDA-approved drugs to abort an acute attack of HAE. However, in resource-poor countries and/or small hospitals or pharmacies where ecallantide, icatibant, or C1-INH replacements are not readily available due to costs or infrequent use, a dose of subcutaneous LMWH should be considered if there are no contraindications in addition to good supportive care in patients with acute exacerbations of HAE (40).

## References

1. Busse PJ, Christiansen SC. Hereditary angioedema. N Engl J Med 2020; 382: 1136–1148.

2. Lee PY, Pearce FL. Histamine secretion from mast cells stimulated with bradykinin. Agents Actions 1990; 30: 67–69.

3. Zuraw BL, Bork K, Binkley KE, Banerji A, Christiansen SC, Castaldo A, Kaplan A, Riedl M, Kirkpatrick C, Magerl M, Drouet C, Cicardi M. Hereditary angioedema with normal C1 inhibitor function: consensus of an international expert panel. Allergy Asthma Proc 2012; 33 Suppl 1: S145–S156.

4. Bafunno V, Firinu D, D’Apolito M, Cordisco G, Loffredo S, Leccese A, Bova M, Barca MP, Santacroce R, Cicardi M, Del Giacco S, Margaglione M. Mutation of the angiopoietin-1 Gene (ANGPT1) Associates With a New Type of Hereditary Angioedema. J Allergy Clin Immunol 2018; 141: 1009–1017.

5. Bork K, Wulff K, Steinmüller-Magin L, Braenne I, Staubach-Renz P, Witzke G, Hardt J. Hereditary Angioedema With a Mutation in the Plasminogen Gene. Allergy 2018; 73: 442–450.

6. Bork K, Wulff K, Rossmann H, Steinmüller-Magin L, Braenne I, Witzke G, Hardt J. Hereditary Angioedema Cosegregating With a Novel Kininogen 1 Gene Mutation Changing the N-terminal Cleavage Site of Bradykinin. Allergy 2019; 74: 2479–2481.

7. Veronez CL, Aabom A, Martin RP, Filippelli-Silva R, Gonçalves RF, Nicolicht P, Mendes AR, Da Silva J, Guilarte M, Grumach AS, Mansour E, Bygum A, Pesquero JB. Genetic Variation of Kallikrein-Kinin System and Related Genes in Patients With Hereditary Angioedema. Front Med (Lausanne) 2019; 6: 19.

8. Cicardi M, Zuraw BL. Angioedema Due to Bradykinin Dysregulation. J Allergy Clin Immunol Pract 2018; 6: 1132–1141.

9. Katelaris CH. Acute Management of Hereditary Angioedema Attacks. Immunol Allergy Clin North Am 2017; 37: 541–556.

10. Lewis LM. Angioedema: Etiology, pathophysiology, current and emerging therapies. J Emerg Med 2013; 45: 789–796.

11. Aygören-Pürsün E, Bygum A, Grivcheva-Panovska V, Mager lM, Graff J, Steiner UC, Fain O, Huissoon A, Kinaciyan T, Farkas H, Lleonart R, Longhurst HJ, Rae W, Triggiani M, Aberer W, Cancian M, Zanichelli A, Smith WB, Baeza ML, Du-Thanh A, Gompels M, Gonzalez-Quevedo T, Greve J, Guilarte M, Katelaris C, Dobo S, Cornpropst M, Clemons D, Fang L, Collis P, Sheridan W, Maurer M, Cicardi M. Oral Plasma Kallikrein Inhibitor for Prophylaxis in Hereditary Angioedema. N Engl J Med 2018; 379: 352–362.

12. Riedl MA, Aygören-Pürsün E, Baker J, Farkas H, Anderson J, Bernstein JA, Bouillet L, Busse P, Manning M, Magerl M, Gompels M, Huissoon AP, Longhurst H, Lumry W, Ritchie B, Shapiro R, Soteres D, Banerji A, Cancian M, Johnston DT, Craig TJ, Launay D, Li HH, Liebhaber M, Nickel T, Offenberger J, Rae W, Schrijvers R, Triggiani M, Wedner HJ, Dobo S, Cornpropst M, Clemons D, Fang L, Collis P, Sheridan WP, Maurer M. Evaluation of avoralstat, an oral kallikrein inhibitor, in a Phase 3 hereditary angioedema prophylaxis trial: The OPuS-2 study. Allergy 2018; 73: 1871–1880.

13. Banerji A, Riedl MA, Bernstein JA, Cicardi M, Longhurst HJ, Zuraw BL, Busse PJ, Anderson J, Magerl M, Martinez-Saguer I, Davis-Lorton M, Zanichelli A, Li HH, Craig T, Jacobs J, Johnston DT, Shapiro R, Yang WH, Lumry WR, Manning ME, Schwartz LB, Shennak M, Soteres D, Zaragoza-Urdaz RH, Gierer S, Smith AM, Tachdjian R, Wedner HJ, Hebert J, Rehman SM, Staubach P, Schranz J, Baptista J, Nothaft W, Maurer M. HELP Investigators. Effect of Lanadelumab Compared With Placebo on Prevention of Hereditary Angioedema Attacks: A Randomized Clinical Trial. JAMA 2018; 320: 2108–2121.

14. Ecker EE, Gross P. Anticomplementary power of heparin. J Infect Dis 1929; 44: 287–295.

15. Poppelaars F, Damman J, de Vrij EL, Burgerhof JG, Saye J, Daha MR, Leuvenink HG, Uknis ME, Seelen MA. New insight into the effects of heparinoids on complement inhibition by C1-inhibitor. Clin Exp Immunol 2016; 184: 378–388.

16. Wijeyewickrema LC, Lameignere E, Hor L, Duncan RC, Shiba T, Travers RJ, Kapopara PR, Lei V, Smith SA, Kim H, James H, Morrissey JH, Pike RN, Conway EM. Polyphosphate is a novel cofactor for regulation of complement by the serpin, C1-inhibitor. Blood 2016; 128: 1766–1776.

17. Wuillemin WA, te Velthuis H, Lubbers YT, de Ruig CP, Eldering E, Hack CE. Potentiation of C1 inhibitor by glycosaminoglycans: dextran sulfate species are effective inhibitors of in vitro complement activation in plasma. J Immunol 1997; 159: 1953–1960.

18. Levine HT, Stechschulte DJ. Possible efficacy nebulized heparin therapy in hereditary angioedema. Immunol Allergy Practice 1992; 14: 31–36.

19. Majluf-Cruz A, Nieto-Martinez S. Long-term follow up analysis of nadroparin for hereditary angioedema: A preliminary report. Int Immunopharmacol 2011; 11: 1127–1132.

20. Weiler JM, Quinn SA, Woodworth GG, Brown DD, Layton TA, Maves KK. Does heparin prophylaxis prevent exacerbations of hereditary angioedema? J Allergy Clin Immunol 2002; 109: 995–1000.

21. Weiler JM, Stechschulte DJ, Levine HT, Edens RE, Maves KK. Inhaled heparin in the treatment of hereditary angioedema. Complement Inflammation 1991; 8: 240–241.

22. Duncan RC, Mohlin F, Taleski D, Coetzer TH, Huntington JA, Payne RJ, Blom AM, Pike RN, Wijeyewickrema LC. Identification of a catalytic exosite for complement component C4 on the serine protease domain of C1s. J Immunol 2012; 189: 2365–2373.

23. Lennick M, Brew SA, Ingham KC. Kinetics of interaction of C1 inhibitor with complement C1s.. Biochemistry 1986; 25: 3890–3898.

24. McRae BJ, Lin TY, Powers JC. Mapping the substrate binding site of human C1r and C1s with peptide thioesters. Development of new sensitive substrates. J Biol Chem 1981; 256: 12362–12366.

25. Eldering E, Huijbregts CC, Lubbers YT, Longstaff C, Hack CE. Characterization of recombinant C1 inhibitor P1 variants. J Biol Chem 1992; 267: 7013–7020.

26. Morrison JF, Walsh CT. The behavior and significance of slow-binding enzyme inhibitors. Adv Enzymol Relat Areas Mol Biol 1988; 61: 201–301.

27. Longhurst H, Cicardi M, Craig T, Bork K, Grattan C, Baker J, Li HH, Reshef A, Bonner J, Bernstein JA, Anderson J, Lumry WR, Farkas H, Katelaris CH, Sussman GL, Jacobs J, Riedl M, Manning ME, Hebert J, Keith PK, Kivity S, Neri S, Levy DS, Baeza ML, Nathan R, Schwartz LB, Caballero T, Yang W, Crisan I, Hernandez MD, Hussain I, Tarzi M, Ritchie B, Králíčková P, Guilarte M, Rehman SM, Banerji A, Gower RG, Bensen-Kennedy D, Edelman J, Feuersenger H, Lawo JP, Machnig T, Pawaskar D, Pragst I, Zuraw BL. COMPACT Investigators. Prevention of Hereditary Angioedema Attacks with a Subcutaneous C1 Inhibitor. N Engl J Med 2017; 376: 1131–1140.

28. Zuraw BL, Cicardi M, Longhurst HJ, Bernstein JA, Li HH, Magerl M, Martinez-Saguer I, Rehman SM, Staubach P, Feuersenger H, Parasrampuria R, Sidhu J, Edelman J, Craig T. Phase II study results of a replacement therapy for hereditary angioedema with subcutaneous C1-inhibitor concentrate. Allergy 2015; 70: 1319–1328.

29. Späth PJ, Wüthrich B, Bütler R. Quantification of C1-inhibitor functional activities by immunodiffusion assay in plasma of patients with hereditary angioedema--evidence of a functionally critical level of C1-inhibitor concentration. Complement Inflammation 1984; 1: 147–159.

30. Hack CE, Relan A, van Amersfoort ES, Cicardi M. Target levels of functional C1-inhibitor in hereditary angioedema. Allergy 2012; 67: 123–130.

31. Caldwell EE, Andreasen AM, Blietz MA, Serrahn JN, VanderNoot V, Park Y, Yu G, Linhardt RJ, Weiler JM. Heparin binding and augmentation of C1 inhibitor activity. Arch Biochem Biophys 1999; 361: 215–222.

32. Caughman GB, Boackle RJ, Vesely J. A postulated mechanism for heparin’s potentiation of C1 inhibitor function. Mol Immunol 1982; 19: 287–295.

33. Hortin GL, Trimpe BL. C1 inhibitor: different mechanisms of reaction with complement component C1 and C1s. Immunol Invest 1991; 20: 75–82.

34. Rent R, Ertel N, Eisenstein R, Gewurz H. Complement activation by interaction of polyanions and polycations. J Immunol 1975; 114: 120–124.

35. Lahiri B, Bagdasarian A, Mitchell B, Talamo RC, Colman RW, Rosenberg RD. Antithrombin-heparin cofactor: an inhibitor of plasma kallikrein. Arch Biochem Biophys 1976; 175: 737–747.

36. Collen D, Semeraro N, Telesforo P, Verstraete M. Inhibition of plasmin by antithrombin-heparin complex. II. During thrombolytic therapy in man. Br J Haematol 1978; 39: 101–110.

37. Colman RW. Hereditary angioedema and heparin therapy. Ann Intern Med 1976; 85: 399–400.

38. Gray E, Mulloy B, Barrowcliffe TW. Heparin and low-molecular-weight heparin. Thromb Haemost 2008; 99: 807–818.

39. Dijk M, Holkers J, Voskamp P, Giannetti BM, Waterreus WJ, van Veen HA, Pannu NS. How Dextran Sulfate Affects C1-inhibitor Activity: A Model for Polysaccharide Potentiation. Structure 2016; 24: 2182–2189.

40. Chan ED, Majluf-Cruz A. Hereditary angioedema (letter). N Engl J Med 2020; 383: e20.

